# Learning to use landmarks for navigation amplifies their representation in retrosplenial cortex

**DOI:** 10.1101/2024.08.18.607457

**Authors:** Lukas F. Fischer, Liane Xu, Keith T. Murray, Mark T. Harnett

## Abstract

Visual landmarks provide powerful reference signals for efficient navigation by altering the activity of spatially tuned neurons, such as place cells, head direction cells, and grid cells. To understand the neural mechanism by which landmarks exert such strong influence, it is necessary to identify how these visual features gain spatial meaning. In this study, we characterized visual landmark representations in mouse retrosplenial cortex (RSC) using chronic two-photon imaging of the same neuronal ensembles over the course of spatial learning. We found a pronounced increase in landmark-referenced activity in RSC neurons that, once established, remained stable across days. Changing behavioral context by uncoupling treadmill motion from visual feedback systematically altered neuronal responses associated with the coherence between visual scene flow speed and self-motion. To explore potential underlying mechanisms, we modeled how burst firing, mediated by supralinear somatodendritic interactions, could efficiently mediate context- and coherence-dependent integration of landmark information. Our results show that visual encoding shifts to landmark-referenced and context-dependent codes as these cues take on spatial meaning during learning.

## Introduction

Precise and reliable spatial navigation is critical for the survival of most mammals and has an accordingly prominent representation in the brain, spread across multiple areas (Vann, Aggleton, and Maguire 2009; Fischer et al. 2020). Spatial navigation is guided by self-localization based on internal movement estimates combined with sensory inputs that allow animals to locate themselves in the environment. Landmarks are sensory cues that can be used to inform an agent about its location within a given spatial context (E Save and Poucet 2000; Etienne et al. 2000; Biro et al. 2007; Julian et al. 2018). For a sensory stimulus to act as a landmark, it must first be associated with spatial meaning (Epstein et al. 2017; Gothard and Skaggs 1996; Chan et al. 2012; Taube and Burton 1995; Jeffery 1998). Once the spatial meaning of a landmark in a given environment has been learned, subsequent exposures to the landmark allow current self-localization estimates to be corrected (Etienne, Maurer, and Séguinot 1996; Campbell et al. 2018; Knierim, Kudrimoti, and McNaughton 1998; Gothard, Skaggs, and McNaughton 1996). How environmental cues are integrated into neural codes of space is an important but poorly understood process that remains a key unanswered question for the field of navigation, and which could shed light on the mechanisms underlying navigational deficits in Alzheimer’s disease and dementia.

Converging evidence points to retrosplenial cortex (RSC) as an important locus for landmark processing (Vann, Aggleton, and Maguire 2009; Etienne, Maurer, and Séguinot 1996; Gothard and Skaggs 1996; Jeffery 1998; Auger, Mullally, and Maguire 2012; Jacob et al. 2017; Fischer et al. 2020; Mao et al. 2020). Neurons in RSC have been shown to encode a range of egocentric and allocentric encoding properties, making it an ideal locus for landmark processing (Alexander and Nitz 2017; 2015; Vedder et al. 2016; Mao et al. 2020; 2017). During spatial navigation, RSC neurons represent visual landmarks via nonlinear integration of self-motion and visual inputs (Fischer et al. 2020). Recordings in freely moving rats have shown that individual RSC neurons conjunctively encode space in egocentric and allocentric spatial reference frames (Alexander and Nitz 2015). Complementary evidence from freely rotating head-fixed mice indicate that top-down dendritic computations are engaged in this process (Voigts and Harnett 2020). These properties suggest that RSC is a key node in processing sensory inputs for effective landmark-mediated self-localization.

However, a physiologically plausible mechanistic understanding of how neurons learn the spatial meaning of landmarks over the course of exploring an environment is missing. Previous studies show an interplay between primary visual cortex and RSC during landmark-dependent navigation (Fischer et al. 2020; Campbell et al. 2018). Other experiments indicate that RSC encodes self-motion information (e.g. how many steps have I taken?), as well as world-referenced inputs (e.g. where in my field of vision is a wall?) (Mao et al. 2020; 2017; Julian et al. 2018). To effectively utilize landmarks, RSC must generate accurate self-localization estimates by combining these two sources of information. Neural networks in RSC are therefore subject to conflicting demands in order to reconcile self-localization estimates with sensory inputs. Internal location representations need to be continuous and resist sudden jumps to provide reliable estimates, even during times of scarce external information, such as low-light conditions (McNaughton et al. 1996; Moser, Kropff, and Moser 2008). Sensory processing, in contrast, must rapidly encode inputs to allow quick responses to novel or unexpected stimuli (D. A. Evans et al. 2018; Carandini and Churchland 2013). How these different coding regimes interact to provide continuous and accurate position estimates is currently poorly understood (T. Evans et al. 2016; Angelaki and Laurens 2020). We hypothesized that neurons in RSC alter the balance of self-refenced or world-reference codes depending on which source of information provides a more accurate self-localization signal in the current environment.

To address this question, we recorded longitudinally from the same RSC neurons as mice learned a landmark-dependent navigation task using 2-photon imaging. We used a generalized linear model (GLM) to evaluate the contribution(s) of self-referenced versus world-referenced factors to the activity of individual neurons over the course of learning. We then assessed neuronal activity in different behavioral contexts to characterize the properties of landmark signals in RSC. Based on our experimental data we developed a proof-of-principle model that uses a putative cellular mechanism for landmark-mediated error correction during navigation. Our results show that individual neurons in RSC shift their activity patterns to allow efficient landmark-mediated self-localization.

## Results

### RSC neurons transition from self-referenced to landmark-referenced spatial codes while learning a visual landmark navigation task

We trained mice to perform a virtual landmark navigation task (Fischer et al. 2020). Briefly, head-fixed mice learned to lick for water rewards at one of two different unmarked reward locations associated with two distinct visual landmarks along a virtual corridor (Fig. 1A, Fig. S1A-C). The virtual corridor consisted of a floor and two walls that contained non-location-specific patterns that provided optic flow but no other spatial information. Each trial started at a randomized location between 50 to 150 cm before a landmark. By licking within the (unindicated) reward zone, mice could trigger water delivery. In-between trials, mice spent at least 3 seconds in a featureless ‘black box.’

**Figure 1:**
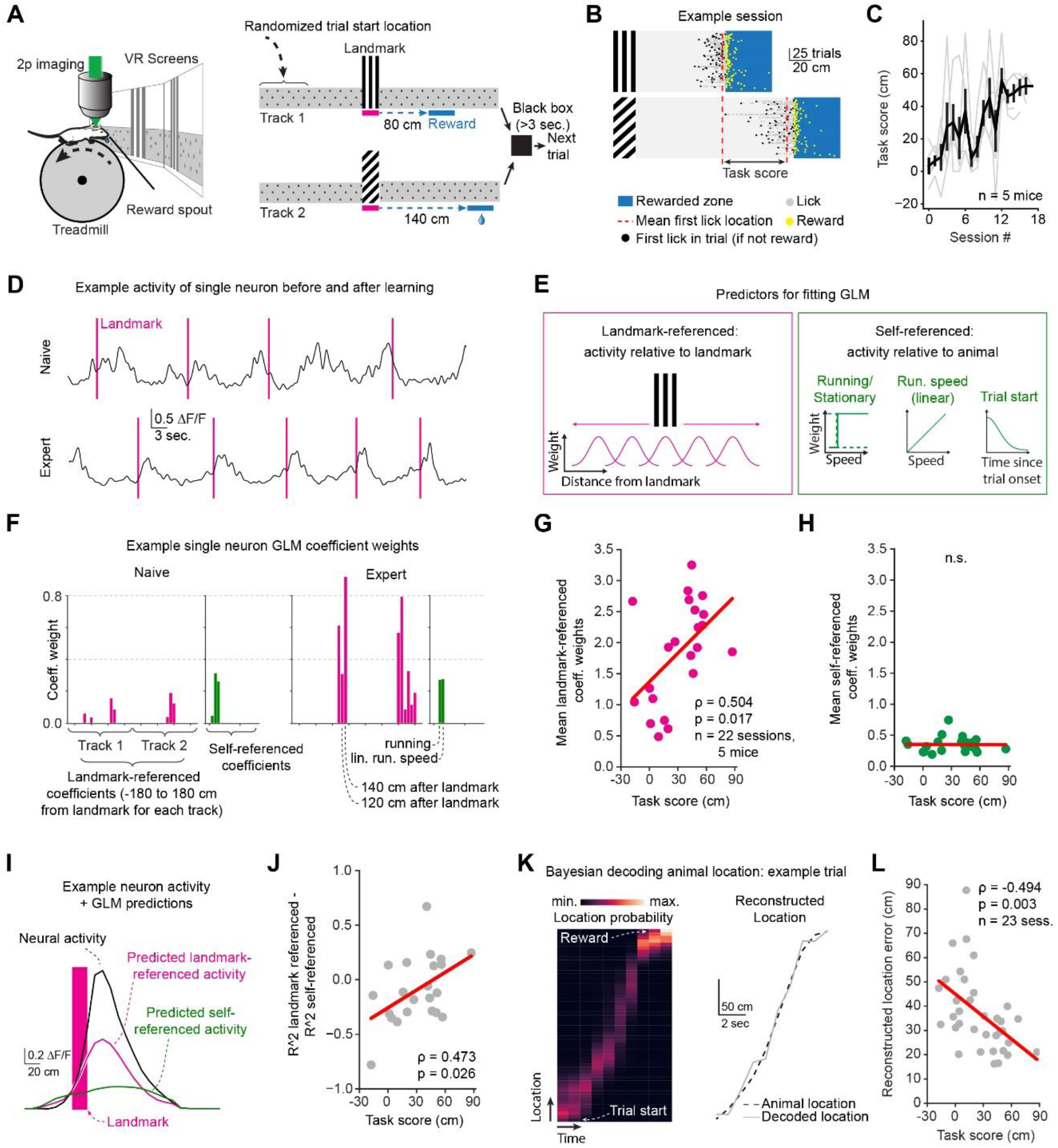
RSC neurons transition from self-referenced to landmark-referenced spatial codes while learning a visual landmark navigation task. (**A**) Experimental setup (left) and task schematic (right). Animals had to traverse a virtual linear corridor and locate an unmarked reward zone relative to one of two visual landmarks that were randomized on a trial-to-trial basis (‘Track 1’ and ‘Track 2’). Animals were then “teleported” to a black box for >3 seconds and subsequently to a randomized start location for the next trial. (**B**) Raster plot of licking behavior relative to the respective landmarks. (**C**) The performance metric (task score) was calculated as the median distance between licking onset on Track 1 and Track 2. (**D**) GCaMP fluorescence of an example neuron over multiple trials before and after learning the task (task score > 20 cm). Purple vertical lines indicate when the animal passed the landmark. (**E**) Predictors used to fit a generalized linear model (GLM). Predictors were categorized into two groups: 1) landmark-referenced (anchored to spatial locations relative to the landmark) and 2) self-referenced to the animal. (**F**) Coefficient weights of the example neuron shown in (D) before and after learning. (**G**) Mean landmark-referenced coefficient weights over the course of learning. **(H)** Same as (G) but for self-referenced coefficients. (**I**) Average activity of an example neuron as a function of location relative to the landmark (black trace) as well as activity predicted by GLM using either landmark (purple) or self-referenced (green) coefficients only. (**J**) Variance explained (R^2^) by prediction of neural activity using only landmark-referenced coefficients and self-referenced coefficients as a function of task score. (**K**) Example trial using a Bayesian decoder to estimate animal position based on neural activity. Left: Probability density function of location estimate for each time bin. Right: Reconstructed location (grey) versus actual location (black). (**L**) Mean position reconstruction error as a function of task score across all animals and sessions.

The mean distance between the first lick location on each track in a given session was used as a behavioral readout for task proficiency (Fig. 1B, C, Fig. S1B). Two-photon GCaMP6f imaging of layer 2/3 (L2/3) RSC neurons was carried out on 35/85 interspersed sessions (n=5 mice) sessions: the other 50/85 sessions were behavior only training sessions (Fig. 1D). A subset of individual neurons were tracked 22 of the 35 sessions (mean ± SEM: 35.14 ± 1.86 tracked cells/session). To do so, we matched the spatial footprints of neurons between a reference session and a second session using the CellReg algorithm (Sheintuch et al. 2017) in conjunction with manual curation (see Methods and Fig. S2A-C).

We applied a generalized linear model (GLM) to quantify which behavioral variables each neuron encoded in each session as animals learned the task. The GLM included two categories of predictors: landmark-referenced and self-referenced. Self-referenced predictors captured behavioral variables relative to the animal, while landmark-referenced predictors related to locations relative to the landmark (Fig. 1E, Fig. S1F). The GLM was fit to the GCaMP fluorescence signals using elastic net regularization (Friedman, Hastie, and Tibshirani 2010) (α=0.5, see Methods for details, Fig. S3A). Coefficient weights showed an increase in the weight of landmark-referenced predictors over the course of learning (Fig. 1F-H, Fig. S1D, E; Pearson correlation ρ=0.504, p=0.017, n=22 sessions, 5 animals, mean ± SEM: 35.1 ± 1.9 tracked neurons/session). These results were in line with previously reported results (Fischer et al. 2020). Further analysis showed that neither the number of non-zero coefficients, peak fluorescence signal, nor trial-by-trial robustness of responses changed over the course of learning. However, neuron activity increased overall (Fig. S3E-H). Predicting neural activity using only landmark-referenced or only self-referenced predictors showed small but significant increases in model fit (explained variance R^2^; Fig. 1I,J, Pearson correlation ρ=0.473, p=0.026), despite the overall explained variance not changing significantly (Fig. S3D) and running speeds remaining largely similar (Fig. S3L). The same relationships held true when all neurons in all recorded sessions, rather than just neurons that were tracked across sessions, were included in this analysis (Fig. S3B, C).

We next evaluated how well RSC neurons encoded space by decoding the animal’s location relative to the landmark using population activity. For this analysis we used all recorded neurons in a given session (mean ± SEM: 75.51 ± 3 neurons/session). We found a significant decrease in reconstruction error as task proficiency increased, suggesting that landmark-anchored spatial codes increase with the animal’s ability to use landmarks for navigation (Fig. 1K, L, Fig. S2I-K; Pearson correlation ρ=-0.494, p=0.003). Together, our results show a significant correlation in RSC neurons between the representation of landmark-referenced activity and proficiency in using landmarks for navigation. This indicates that RSC neurons shift their encoding priorities as a function of task demands.

### Visual inputs stabilize spatial codes in RSC

We next asked how stable landmark representations in RSC are over time. Persistent codes allow for stable output to downstream structures, while variable population activity indicates that organizational principles are embedded in higher-order network dynamics (Quian Quiroga and Panzeri 2009; Haider et al. 2016; Ujfalussy et al. 2015; Remington et al. 2018). Previous work has shown that spatial codes of individual RSC neurons vary day-by-day in the absence of location-specific visual inputs (Mao et al. 2018). We therefore asked whether learning visual landmark cues can stabilize neuronal representations across days. We tested the persistence of landmark-referenced codes in RSC by tracking the same neurons over three expert sessions (task score > 20 cm; Fig. 2A & Fig. S2A-C). The task score threshold of 20 cm was empirically chosen to indicate when mice began reliably using landmarks to locate rewards. These sessions constitute a subset of the sessions shown in Fig. 1. Peak responses (calculated as the peak average ΔF/F across all trials along the track) and the area under the curve of all tracked neurons (n = 169 neurons from 5 animals) did not change significantly across days (Fig. 2B, C; One-way ANOVA, p=0.5 for ΔF/F peak, p = 0.8 for AUC/spatial bin).

**Figure 2:**
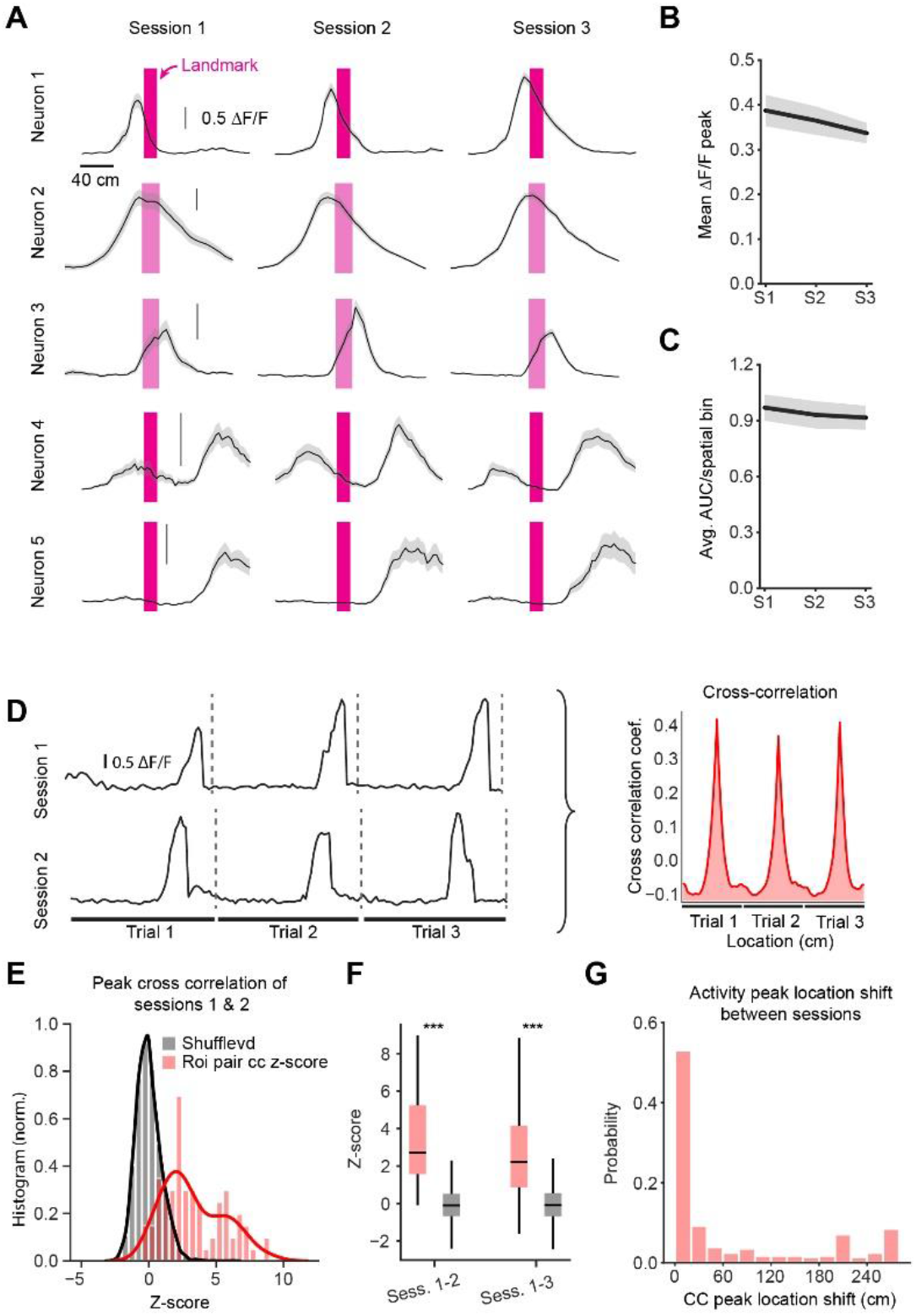
RSC neuron spatial codes stabilize after learning in the presence of visual cues. (**A**) Trial-averaged activity of five example neurons as a function of location on the track. All vertical scale bars indicate 0.5 ΔF/F. (**B**) Mean peak ΔF/F for all tracked neurons across three sessions. (**C**) Same as (B) but for area under the curve (AUC) calculation. One-way ANOVA indicates no significant difference between days. (**D**) Schematic of cross-correlation calculation between sessions. For each neuron, the cross correlation between sessions was calculated from spatially binned activity with periods in-between trials removed. (**E**) Histogram of the z-score of each neuron’s cross-correlation value relative to a shuffled distribution. (**F**) Z-score boxplot for first vs. second and first vs. third session. (**G**) Probability of cross-correlation peak location shift.

To test if neurons retained their spatial tuning, we quantified the cross correlation of each neuron’s activity as a function of location on the track. The cross-day cross-correlation was calculated by concatenating the activity of a given neuron on all trials where a given landmark was shown and finding the highest peak in the cross-correlogram (Fig. 2D). Neural activity in the black box between trials and after reward delivery was removed from this analysis. Neurons with an R^2^ value smaller than 0.25 were not included in this analysis to remove neurons with little-to-no landmark-or self-referenced activity. In total, 156 neurons (out of 169 tracked neurons) from 5 mice were used in this analysis. To test whether individual neurons were significantly cross-correlated across days, a shuffled distribution of cross-correlation values was calculated for each neuron by rotating its neural activity by a random amount for each trial and re-calculating the cross-correlation. We used this shuffled null distribution to compare trial-by-trial activity correlations of individual neurons to spatially-randomized activity. The z-score relative to the shuffled distribution was then calculated (Fig. 2E). We found that landmark-referenced codes across three consecutive sessions were stable (Fig. 2F; median z-score for correlation between sessions 1-2: 2.72, IQR: 3.67; session 1-3: 2.21, IQR: 3.29, unpaired two-tailed T-test: p<0.0001 for all comparisons, see Fig. S4A for non-z-scored cross correlation values). Finally, we analyzed the peak-shift in cross correlation (Fig. 2G). The peak-shift analysis revealed a median shift of 15 cm (IQR: 135 cm), suggesting that the vast majority of neurons retain their spatial tuning. These results indicate that behaviorally-relevant anchoring visual cues can stabilize spatial codes in the cortex.

### RSC neurons are most active when visual flow and self-motion signals are coherent

Context-dependent processing of sensory inputs is a cornerstone of cortical function (Mante et al. 2013; Smith, Barredo, and Mizumori 2012; Mao et al. 2017). A number of studies have reported a general decrease in activity and spatial specificity during visuo-motor mismatch conditions in cortex (Harvey, Coen, and Tank 2012; Fischer et al. 2020; Diamanti et al. 2019; Mao et al. 2020). However, a more detailed understanding of the patterns of activity changes can provide clues as to how visual and motor information is integrated. We hypothesized that RSC neurons implement context-sensitive codes by associating self-motion with visual motion cues in a given environment. This allows sensory cues to be differentially encoded depending on the behavioral context in which they are encountered. To test this hypothesis, we introduced visuo-motor mismatch trials in which virtual movement was uncoupled from animal running (Fig. 3A). During these mismatch trials, the virtual corridor moved at a constant speed of 30 cm/sec, independent of running behavior. The reward zone was in the same virtual location and, though licking behavior was still recorded, no rewards were dispensed. A brief trial break (approx. 30 sec.) between VR and visuo-motor mismatch sessions signaled the behavioral context switch. Mice ceased to lick in the rewarded zone soon after the switch to the visuo-motor mismatch session (Fig. 3B), behavioral confirmation that they recognized the change. Congruent with previous data (Fischer et al. 2020), neural activity overall was lower during the visuo-motor mismatch session compared to when the animal was actively executing the task (Fig. 3C; mean ± SEM VR: 0.31 ± 0.03, visuo-motor mismatch: 0.19 ± 0.02, paired, two-tailed T-test: p=0.001, n=13 sessions, 5 naïve and 8 expert with a task score >20 cm, n=5 mice).

**Figure 3:**
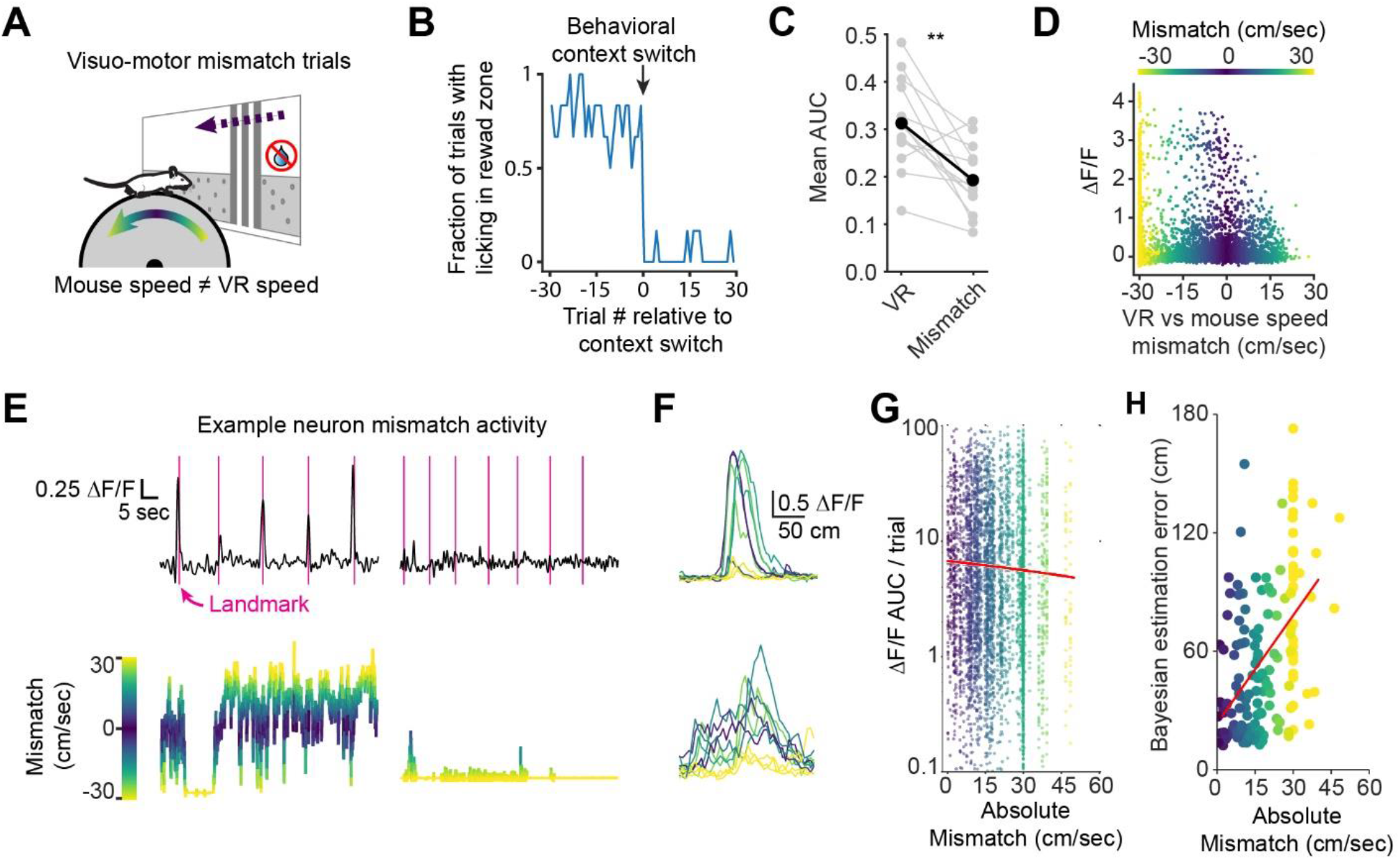
Behavioral context modulates RSC neuronal response strength and spatial coding. (**A**) Visuo-motor mismatch experiment schematic: virtual corridor flow speed (30 cm/sec) versus animal running speed was measured in relation to neuronal responses. (**B**) Fraction of trials with at least one lick in the reward zone. (**C**) Mean GCaMP6f ΔF/F signal area under the curve (AUC) per 5 cm spatial bin during VR and visuo-motor mismatch periods. (**D**) ΔF/F of an example neuron as a function of mismatch. Each dot represents the signal of one recorded frame. Color code indicates speed mismatch. (**E**) Example GCaMP6f ΔF/F traces of one neuron (top) and the difference in animal running and virtual corridor flow speed (ΔSpeed; below) during two sets of trials (left: 5 trials where the animal was locomoting, right: 7 trials where the animal was quiescent). (**F**) GCaMP6f activity of two example neurons during visuo-motor mismatch session. Each trace represents one trial with the color code indicating the difference in average running and virtual corridor flow speed for that trial. (**G**) Integrated ΔF/F of each trial as a function of speed mismatch for all neurons and all trials of n=8 expert sessions. Each trial of a given neuron is represented by one dot. Red line: linear fit. (**H**) Location reconstruction error using a Bayesian decoder that was trained on virtual navigation data and applied to visuo-motor mismatch trials (n=8 expert sessions). Red line: linear fit.

We analyzed neural activity in visuo-motor mismatch sessions as a function of the difference between the visual flow speed and the animal’s running speed on the treadmill in expert animals (Fig. 3D). To account for linear speed modulation of neural activity, we fit a linear regression to the activity of each neuron as a function of speed and subtracted the corresponding linear factor (see Methods). We found a tendency for neurons to be most strongly activated when the movement speed of the displayed corridor matched the animal’s own running speed (Fig. 3E,F, Fig. S5A). To quantify the relationship between visuo-motor mismatch and neural activity, we calculated the integrated calcium activity of each neuron on each trial as a function of the absolute visuo-motor mismatch. This revealed a modest negative correlation, suggesting that neurons are most active when visuo-motor mismatch was smallest (Fig. 3G, Spearman rank correlation: ρ=-0.036, p=0.0017; the same relationship held true when we did not correct for linear speed, Fig. S5C). No relationship between running speed and neural activity was found while animals were in the black box in-between trials (Fig. S5B). We reasoned that the relationship between mismatch and neural activity should translate into improved spatial coding when running and visual flow speed match, and vice versa, if the mismatch is large, spatial coding should be disrupted. To test this, we trained a Bayesian decoder on neural data collected during virtual navigation in trained animals (task score > 20 cm) and used it to estimate the mouse’s location during visuo-motor mismatch trials. Reconstruction error was smallest when the difference in animal running speed and visual flow speed was around zero (Fig. 3H, Spearman rank correlation: ρ=-0.474, p<0.001). These results indicate that RSC neurons are functionally organized to most strongly represent visual feedback when coordinated with self-motion, thus providing evidence for behavioral-context representation that is based on reconciling internal and external cues for self-localization.

### Bursting firing mediated by dendrites can accurately correct self-localization estimates

Our experimental results show that RSC neurons develop stable, landmark-referenced codes over learning that are modulated by behavioral context. We hypothesized that individual neurons could generate these landmark codes by integrating visual bottom-up inputs with contextual top-down signals. Recent work has shown that RSC receives spatially segregated inputs from thalamic, primary visual, and associative cortices (Lafourcade et al. 2022). We reasoned that the generation of bursts of action potentials generated by coincident bottom-up somatic and top-down dendritic inputs (Naud and Sprekeler 2018; Payeur et al. 2021; Xu et al. 2012; Ranganathan et al. 2018; Bicknell and Häusser 2021; Takahashi et al. 2016; Francioni and Harnett 2021; London and Häusser 2005; Larkum, Kaiser, and Sakmann 1999; Larkum 2013; Greedy et al. 2022; Fişek et al. 2023) could underlie the amplification of sensory inputs by behavioral context we have observed in our data. We therefore created a model to explore how somatodendritic interactions in pyramidal neurons could potentially be utilized to provide context-dependent self-localization.

Our model combined multi-compartment spiking neurons developed by Naud and Sprekeler (2018) with an attractor model representing an agent’s self-localization estimate (Ocko et al. 2018). We adjusted the parameters of neurons compared to (Naud and Sprekeler 2018), which consisted of a somatic and a dendritic compartment (Fig. 4A,B). The biophysical properties of each compartment endowed these neurons with the ability to generate bursts of action potentials, but only when somatic and dendritic inputs coincided (Fig. 4B,C) (Naud and Sprekeler 2018). The multi-compartment neurons sent output to a linear attractor (Fig. 4A, Fig. S6A), which represented the agent’s self-localization estimate by a Gaussian activity bump centered at the best current location estimate (Zhang 1996; Campbell et al. 2018; Ocko et al. 2018). Each neuron received visual inputs at a location relative to the landmark. The location relative to the landmark was drawn from a Gaussian distribution (Fig. S8A). Even though visual landmark inputs cease after the animal has passed the landmark, in this model some visual inputs indeed have their receptive field center after the landmark. This is consistent with previous findings (Fischer et al. 2020) and is further supported by the rich responses primary visual cortex is known to generate (Saleem et al. 2018; Niell and Stryker 2010; Musall et al. 2019; Pakan et al. 2018). Visual landmark inputs were modeled as somatic input currents that scaled as a function of distance to its respective receptive field center.

**Figure 4:**
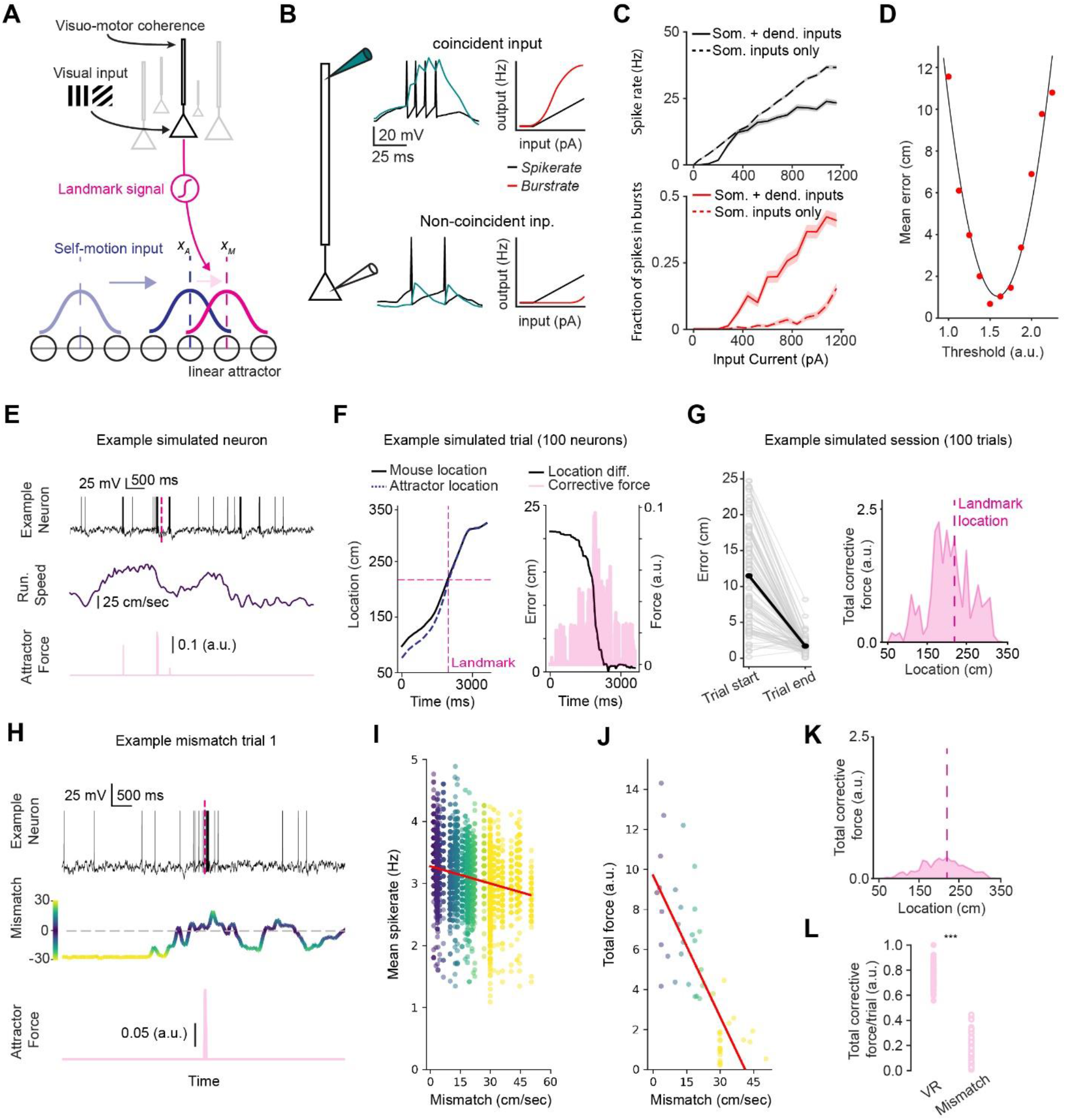
Somatodendritic bursting in a hybrid simulated neural network enables accurate location error correction. (**A**) Simulation components. Each landmark neuron is connected onto the attractor at its respective spatial receptive field center. (**B**) Schematic of spikes and bursts generated as a function of somatic and dendritic inputs. Bursts are defined as 2 or more spikes with an interspike interval <10 ms. (**C**) Spike and burst rates in response to increasing current inputs. (**D**) Agent self-localization error correction during simulated trials as a function of threshold between neurons and attractor. Each red dot represents a set of simulated trials at a given threshold value, black line is a 2nd order polynomial fit. (**E**) Example simulated trial. Top: example neuron activity. The landmark is indicated by the dashed lines. Middle: running speed in simulated trial. Bottom: force applied to attractor by example neuron. (**F**) Left: Actual location (solid line) and self-localization estimate of attractor (dashed line). Right: Self-localization error (calculated as actual location minus attractor activity bump location) and corrective force applied to the attractor by 100 landmark neurons. (**G**) Left: Error at the beginning and at the end of 100 simulated trials. Right: Total force applied to the attractor by all landmark neurons combined. (**H**) Activity traces of an example neuron during visuo-motor mismatch. Neural activity is shown above the respective speed mismatch (color coded) and the force exerted by that neuron on the attractor (pink). (**I**) Mean spike rate per trial as a function of visuo-motor mismatch speed. Each dot represents one trial of one neuron. (**J**) Force exerted on attractor. (**K**) Total force exerted by landmark neurons on the attractor during mismatch trial simulation. (**L**) Force exerted onto attractor during simulated virtual navigation (“VR”) and visuo-motor mismatch trials (“Mismatch”).

Each neuron also received input into its dendritic compartment as a function of coherence between self-motion and visual flow feedback (visuo-motor coherence; Fig. 4A). Both compartments received constant noise inputs (see Methods) that resulted in stochastic background spiking. The combination of bottom-up feedforward and top-down feedback inputs resulted in an increased propensity for neurons to burst. In contrast, neurons receiving an equivalent total amount of current injected into the somatic compartment alone emitted significantly fewer bursts (Fig. S6B). Each spike emitted by a landmark neuron exerted a rapidly decaying force on the attractor via a sigmoidal activation function that incorporated a gating threshold (Fig. 4A; Methods) that corrected the self-localization estimate towards its respective receptive field center. This rapid decay acted as a filter that prevented the self-localization estimate from erroneously changing due to background noise. To test this, we ran a series of simulations with increasing gating threshold values (Fig. 4D). While a low threshold value meant single spikes pull the attractor away from the agent’s actual location, higher thresholds prevented any error correction altogether. Our fit showed an optimal range between 1.4 and 1.8. We used 1.75 for all subsequent simulations.

We then used behavioral data from our experimental recording sessions to simulate 100 trials of the landmark navigation task (from Fig. 1) with 100 simulated neurons whose activity was anchored by the landmark (“landmark neurons”). Each trial started with a random initial offset between the self-localization estimate and the agent’s actual location. As the agent traversed the linear corridor, the combined force generated by neurons corrected the self-localization estimate (Fig. 4E-G, Fig. S8B; mean final error: 1.7 ± 0.11 cm. Paired t-test: p < 0.001; mean corrective force AUC ± SEM: 225.11 ± 2.62). In contrast, running the same simulation without coincident somatodendritic inputs resulted in significantly worse error correction (Fig. S6C-E, Fig. S8C). These simulations show that bursts of actional potentials generated by somatodendritic interactions are a robust way to generate corrective inputs for self-localization estimates.

### Somatodendritic interactions can support context-dependent computations

Our experimental data indicate that mouse RSC exhibits different sensory cue integration regimes during virtual navigation and visuo-motor mismatch sessions. We tested if our model could recapitulate these findings by simulating 100 visuo-motor mismatch trials. Consistent with our virtual navigation simulation, we used behavioral data from our mouse experiments. We reasoned that visual landmark inputs do not influence downstream neural representations during visuo-motor mismatch, as they are not relevant to ongoing behavior, and they should therefore exert minimal force. We modeled dendritic inputs as a visuo-motor coherence signal which was dependent on the difference in agent running and VR flow speed. The virtual environment was shown at a constant flow speed of 30 cm/sec while mice were free to sit or run on the treadmill. Similar to observations made in biological neurons during this task (Fig. 3), the activity of model neurons was highest when animal running speed approximately matched visual flow speed leading to increased force exerted on the attractor (Fig. 4H-J, Spearman rank correlation for spike rate: ρ=-0.263, p<0.001, for force exerted: p=-0.799, p<0.001). Overall, the total force exerted on the attractor was significantly lower compared to virtual navigation during visuo-motor mismatch trials (Fig. 4K,L; mean ± SEM VR: 225.11 ± 2.62, Mismatch: 42.25 ± 3.43, Two-tailed T-Test for related samples: p < 0.001). By modeling dendritic inputs as visuo-motor coherence, we were thus able to implement a biophysically-plausible mechanism for context-dependent sensory cue integration switching.

Finally, we tested if supralinear somatodendritic interactions in our model were necessary for context-dependent landmark computations. We ran two simulations using linearly or supralinearly integrating single-compartment variants of the previously described artificial neurons (Fig. S6A,B). Both models had an otherwise identical architecture. Supralinear single-compartment neurons transformed current input into spiking in such a way that it mirrored that of 2-compartment neurons with coincident somatodendritic inputs (Fig. S6C, top). This is achieved by multiplying input currents by a factor that is dependent on the membrane potential (see Methods). Vice-versa, linear single-compartment neurons matched the firing properties of 2-compartment neurons without coincident inputs (Fig. S6C, bottom). The supralinear model reliably corrected the agent’s self-localization estimate (Fig. S6D). In contrast, the linear model did not (Fig. S6E). The lack of correction in the linearly integrating model is the result of the landmark neuron’s inability to exert any meaningful force on the attractor (Fig. S6F; mean corrective force AUC ± SEM supralinear model: 711.05 ± 11.63; linear model: 0.001 ± 0.0008). This is reflected in the overall correction performance over 100 trials (Fig. S6G). Importantly, neither model was able to respond appropriately to the behavioral context switch. The supralinear single-compartment model fired bursts consistently, regardless of behavioral context (Fig. S6H; Fig. S9A,B; mean corrective force AUC ± SEM during VR: 711.05 ± 11.63; during visuo-motor mismatch: 225.11 ± 2.62). In contrast, the linear model was never able to correct the agent’s self-localization estimate (Fig. S6I; Fig. S9C,D; mean corrective force AUC ± SEM during VR: 0.001 ± 0.0008; during visuo-motor mismatch: 2.59 ± 4.3×10-6). Together, these results provide evidence that supralinear dendritic integration in 2-compartment cortical neurons can facilitate cortical computations across behavioral contexts (Tran-Van-Minh et al. 2015; Francioni and Harnett 2021; Poirazi and Papoutsi 2020; Greedy et al. 2022; B. A. Richards and Lillicrap 2019; Payeur et al. 2021).

## Discussion

We provide evidence for a learned, persistent, and context-dependent landmark code in RSC. This code is evident in a significant increase in the encoding of landmark-referenced variables over the course of task acquisition (Fig. 1) that remains stable over days (Fig. 2). The landmark codes are significantly attenuated when landmarks are irrelevant to behavior, but appear to increase activity if external visual cues and animal behavior are similar to the context in which landmark signals are important (Fig. 3). Based on our data, we combined a multi-compartmental neuron model with an attractor network to evaluate the plausibility of a somatodendritic mechanism in which burst firing of cortical neurons mediates a corrective signal.

While numerous studies have shown that RSC is involved in landmark processing (Fischer et al. 2020; Epstein 2008; Maguire 2001), how individual neurons adapt their encoding properties while learning the spatial meaning of a landmark is unknown. We show that individual RSC neurons significantly increase their encoding self-referenced variables over the course of learning. Interestingly, our GLM results further indicate that neurons retain some encoding of self-referenced information. This result is consistent with the demands of our task, which requires animals to use landmark-referenced information during self-localization as well as self-referenced information during the localization of the reward after they have passed the landmark. Our results are congruent with previous studies that point to conjunctive encoding of ego- and allocentric variables in RSC (Alexander and Nitz 2015; Mao et al. 2020; Jacob et al. 2017). It is worth noting that residual encoding of self-referenced variables may account for path-integration deficits found in lesion studies of RSC (Cooper and Mizumori 2001; 1999; Elduayen and Save 2014). An exciting direction for future inquiries is to gain a deeper understanding of how task structure affects encoding priorities in RSC.

Landmark codes were stable across days in our paradigm. This contrasts with previous observations of variation across days in the spatial tuning of RSC cells (Mao et al. 2018). We posit that the presence of visual landmarks in our experimental design underlies the cross-day stability that we have observed. This result could be related to instability of other cell types that are tuned either to the environment or oneself, such as place cells (Etienne Save, Nerad, and Poucet 2000; Muller and Kubie 1987), head-direction cells (Taube, Muller, and Ranck 1990; Knight et al. 2014) and even grid cells (Campbell et al. 2018). This speaks to the importance of visual landmarks in spatial navigation. However, the exact topological organization of spatially tuned brain structures, and how visual inputs are integrated during navigation, remains unknown. Our work suggests that RSC is a key node in receiving and parsing behaviorally relevant visual stimuli during navigation.

Context-dependent modulation of neural activity is a critical aspect of cortical computation and has been found in nearly every region where it has been investigated (Harris and Mrsic-Flogel 2013; Pakan et al. 2016; Zipser, Lamme, and Schiller 1996; Ferguson and Cardin 2020). In line with previous reports in RSC (Fischer et al. 2020; Mao et al. 2017; 2020; Harvey, Coen, and Tank 2012), we found a decrease in overall neural activity when animals are not actively navigating. Such an overall decrease could be attributed to a number of factors including overall decreased engagement or lack of reward. While we cannot entirely rule out other contributing factors, our previous work (Fischer et al. 2020) and our analyses here suggest that general attenuation of neural activity is not the most likely explanation. Our data further showed that neuronal activity increases as the mismatch between visual flow and self-motion feedback decreases. One possible mechanism underlying such a context-dependent modulation could be predictive coding, which may be dendrite-mediated (Rao and Ballard 1999; Leinweber et al. 2017; Keller and Mrsic-flogel 2018). While previous studies investigating dendritic mechanisms of nonlinear integration have mostly focused on primary sensory areas (Xu et al. 2012; Ranganathan et al. 2018; Francioni, Padamsey, and Rochefort 2019; Beaulieu-Laroche et al. 2019; Palmer et al. 2014; Manita et al. 2015; Ayaz et al. 2019), associative areas may implement similar computations (Lafourcade et al. 2022).

We have posited that visuo-motor coherence could provide a top-down signal for when a cue is encountered in a familiar location. This enables correct encoding of landmarks in two ways: 1) A visual cue passes through the visual field at different rates, depending on its distance from the observer. Visuo-motor coherence thus allows correct encoding of visual cue distance; 2) When a cue is encountered but the animal is not actively navigating, as is the case in mismatch sessions, the same cue does not generate bursts and thus does not affect downstream change in positional codes. Our simulations predict a dendrite-localized signal that varies as a function of the learned association between a given, familiar environment and self-motion. Future studies may address this mechanistic hypothesis resulting from our work.

We simulated a population of multi-compartment neurons (Naud and Sprekeler 2018) to explore how the computational capabilities of cortical neurons add flexibility to the way visual landmark inputs update self-localization estimates. Previous studies have investigated how corrective inputs to attractor networks can offset errors that either accumulate over time or are introduced through environmental manipulations (Campbell et al. 2018; Hardcastle, Ganguli, and Giocomo 2015; Bicanski and Burgess 2016; Burak and Fiete 2009; Page and Jeffery 2018). Most of these models are designed to work in a single behavioral context and are therefore unable to account for more complex demands in real world scenarios. We explicitly modeled the source of landmark and context signals to show how neuronal output can be efficiently modulated by a simple somatodendritic mechanism. We note that our experimental paradigm does not capture the complexities of freely moving mice in their natural habitats. However, we contend that our proposed mechanism can generalize to natural environments in which a given landmark, encountered from the same viewing angle under similar self-motion aspects, should still elicit a stronger response compared to the same landmark being seen from completely different vantage point, as self-localization errors can be most efficiently corrected if a landmark is seen from a familiar location.

An important future avenue of inquiry will be to investigate how individual neurons bind their code to a certain location relative to the landmark. A number of studies have looked into this dynamic (Widloski and Fiete 2014; Ocko et al. 2018; Campbell et al. 2018). Our previous work (Fischer et al. 2020) has shown that primary visual cortex sends spatially tuned inputs to RSC which could act as an initially context-free feedforward/bottom-up signal. Over the course of learning, top-down context signals, such as we suggest in this study, could trigger AP bursts which in turn strengthen synaptic connections between RSC neurons and downstream brain structures. Future work is needed to illuminate how burst-dependent synaptic plasticity can play the dual role of controlling synaptic plasticity (Payeur et al. 2021; Greedy et al. 2022; N. A. Richards et al. 2019) while also acting as a form of efficient, context-depending communication between brain structures as we show here (Krahe and Gabbiani 2004; Bialek et al. 1991; Bair et al. 1994).

Investigating dendritic computation in awake behaving animals is currently limited by a number of serious technical and experimental challenges (Francioni and Harnett 2021). The modeling approach we used here has successfully demonstrated complex encoding schemes in the past (Kaifosh and Losonczy 2016; Williams et al. 2021; Payeur et al. 2021; Bicknell and Häusser 2021) and may prove an advantageous complementary avenue to investigate the contribution of dendritic computations during behavior.

Overall, we demonstrate how individual neurons shift their encoding priorities from self-centered to world-centered in our landmark navigation task. These codes remain stable while animals repeatedly execute the same task over multiple days but significantly change their activity patterns in a different behavioral context. However, when behavioral and sensory inputs matched in this alternative context, we observed activity reminiscent of that recorded during active navigation. We formulated a mechanistic hypothesis of how sensory information could be integrated by multi-compartment cortical neurons to update downstream internal state representations through bursting. These bursts were critical for output to downstream neurons while being robust to background noise. Our proposed mechanism therefore combines the advantages afforded by rapid and flexible integration sensory stimuli with the robustness of attractor dynamics for the internal representation of behavioral state.

## Methods

### Animals and surgeries

All animal procedures were carried out in accordance with NIH and Massachusetts Institute of Technology Committee on Animal care guidelines. Male and female mice were singly housed on a 12/12 hr (lights on at 7 am) cycle. Surgical procedures where identical to those in Fischer et.al. 2020. Briefly, C57BL/6 mice aged 7-10 weeks were anaesthetized. A 3.0 mm diameter craniotomy was drilled on the dorsal surface of the skull. AAV1.Syn.GCaMP6f.WPRE.SV40 was injected into the exposed retrosplenial cortex at 4-6 injection sites 1800 -3400 µm caudal of bregma and 350-600 µm lateral to the midline. Recordings were taken directly over the injection sites adjacent to the central sinus. 50-100 nl were injected at each site. After successful injection, a cranial window was placed over RSC and a headplate implanted on the skull.

### Virtual reality setup and behavioral training

The same virtual reality setup as described in Fischer et.al. 2020 was used for this study. Mice were head fixed atop a 20 cm polystyrene disc. Two 23.8” computer screens covered the majority of the mouse’s field of view. During virtual navigation, animal movement was translated into visual flow through the virtual environment that was shown on the screens. A lick spout was placed close to the mouse’s mouth such that it could easily touch it by extending its tongue. Recordings were obtained from mouse from day 0 of exposure to the landmark navigation task. Prior to behavioral training/recording mice were habituated to head-restraint on the treadmill and with a linear corridor without landmarks.

### Image Registration, ROI detection and ROI matching across days

Two-photon imaging was carried as described in Fischer et.al. 2020. In brief, a Neurolabware 2-photon microscope with a 16x objective was used to collect all imaging data. Images were acquired either at 15.5 or 31.0 Hz using am excitation wavelength between 920 and 980 nm. Frames of the raw video data were registered, and putative neurons (regions of interest or ROIs) were detected using the CaImAn software package (Giovannucci et al. 2019). ROIs were subsequently manually curated to remove dendrites or other, non-soma ROIs. Individual neurons from a subset of recorded fields of view (FOV) were tracked across sessions to analyze their activity patterns over the course of learning. In order to do so, one FOV from each animal after it had robustly learned the task (Task score > 20 cm) was used as a reference session. The mask of detected ROIs for any two sessions was then overlaid and coarsely aligned if necessary. This was followed by applying the CellReg algorithm to determine if two ROIs are the same neuron base on centroid distance and spatial footprint (Sheintuch et al. 2017). Supplementary figure 2 shows this process for three FOVs. This process was repeated until each session of a given animal was matched to the reference session. Neurons considered ‘tracked neurons’ are those that have been matched between any given session and the reference session. For the analyses shown in figure 2, the same neurons were tracked over three sessions. This was done by overlaying the ROI masks and selecting ROIs that could be matched across all three sessions.

### GLM

We used a generalized linear model to identify which variables each neuron was encoding throughout training. We created two broad categories of predictors: self-centered and landmark-centered. Self-centered predictors consisted of: Running speed (linear) and categorical (running/not running) with a threshold of 1 cm/sec. Trials starts were captured by 3 Gaussians (standard deviation of 0.25 sec.) offset from the trial start to capture neural activity that is related to the trial start but delayed in terms of fluorescence increase. Landmark-referenced predictors consisted of a series of Gaussians covering the entire virtual corridor. Each landmark had its own set of landmark-referenced predictors. We fit the model on calcium fluorescent data using elastic net regression using the glmnet package (Friedman, Hastie, and Tibshirani 2010) with α=0.5. Only neurons that our model fit reasonably well (explained variance > 25%) were included for GLM analyses in Fig. 1. For cross-validation we held out 1 trial and fit to the others. We repeated this until each trial was held out once. We calculated an R^2^ value (explained variance) to evaluate the fit quality of our GLM model by calculating the difference between the predicted and true calcium signal on held out data:

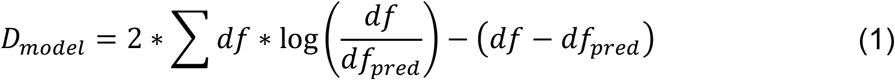

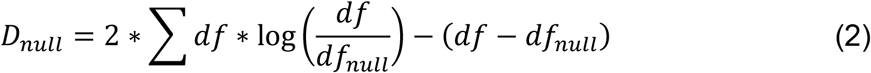

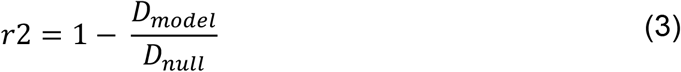

Where df is the average activity of the neuron on the held out trial, df_pred_ is the predicted activity and df_null_ is the average activity of that neuron on that trial. R^2^ was calculated only on activity in-between trial start and reward delivery. We determined the fraction of landmark-referenced vs. self-referenced encoding by calculating the sum of all absolute landmark-referenced coefficient weights divided by the sum of absolute landmark-referenced and self-referenced coefficient weights.

We quantified the explained variance of landmark-referenced of self-referenced coefficients by first fitting the full model but predict activity with either one or the other set of predictors set to 0. This way, only activity captured by the respective coefficients was used for the prediction. Explained variance was calculated as described above.

### Bayesian location decoding

We used a Bayesian decoder to estimate the animal’s position on the track based on neural data alone (Davidson, Kloosterman, and Wilson 2009; Mao et al. 2018). Briefly, spatial tuning curves for each neuron were constructed using one of two methods: 1) In experiments where we analyzed precision of the spatial code during virtual navigation (Fig. 1) every other trial in a session was used to construct the tuning curves. Activity on the other half of trials was used to decode animal position. 2) In experiments that analyzed how well we could reconstruct animal position during visuo-motor mismatch trials, all trials during virtual navigation were used to construct tuning curves while trials during visuo-motor mismatch were used for location estimation.

Location and activity data were binned into 2.5 cm and 0.25 sec. bins, respectively. For each time bin, the probability density function across all location bins on the track was calculated. The reconstructed position was defined as the location with the highest probability in a given time bin. The overall decoding error for a trial was calculated as the median difference between the decoded location and the animal’s actual location. The probability density functions were calculated as described in (Mao et al. 2018):

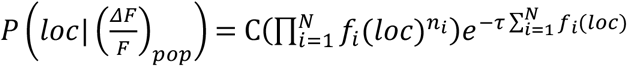

Where *f*_*i*_ is the spatial tuning for neuron *i, N i*s the total number of neurons, *n*_*i*_ is the activity of neuron *i* at the respective time bin (calculated as the average value of all datapoints within that time bin) and *τ* is the time bin size.

### Cross-day cross-correlation analysis

To evaluate the stability of landmark-referenced codes we cross correlated activity over subsequent recording sessions. To do so, we excised neural activity in the black-box between trials and concatenated the remaining calcium fluorescence trace. The resulting traces, one from each recording session, where then shifted relative to each other with a maximum lag of 800 seconds. The maximum cross correlation value was used for further analysis. To create null-distributions for each neuron, we randomly rotated neural activity in one of the sessions for each trial independently. The cross correlation for the null distribution was thus calculated from a non-rotated first session, and a rotated second session.

### Visuo-motor mismatch analysis

We calculated visuo-motor mismatch by subtracting the visual flow speed of 30 cm/sec from the animal running speed. Each visuo-motor mismatch session consisted of 10 cm/sec and 30 cm/sec flow speed sessions. However, here we only used 30 cm/sec trials as these were closer to the animals average running speed and thus gave us better sampling of a range above and below that speed. Before relating flow speed to neural activity, we removed the linear speed component. We did this by fitting a robust linear regression (scipy.stats.siegelslopes) to fluorescence data as a function of running speed. We then subtracted the corresponding amount from each datapoint. For the following analysis we calculated the average running speed on a trial and subtracted the visual flow speed to determine ΔSpeed. To determine the maximum response across all neurons and trials, we evaluated dF/F for each trial as a function of ΔSpeed and fit a 2^nd^ order polynomial (numpy.polyfit).

### 2-compartment model

Variables and parameters are defined in the tables below.

Landmark cells are simulated with a two-compartment model developed by Naud and Sprekeler 2018). The dynamics of the somatic compartment are

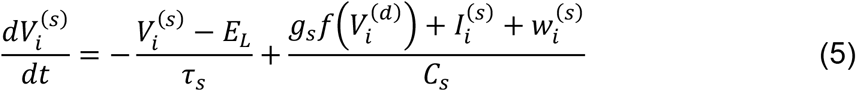

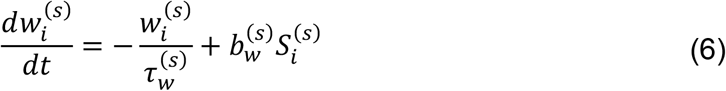

whereas the dynamics of the dendritic compartment are

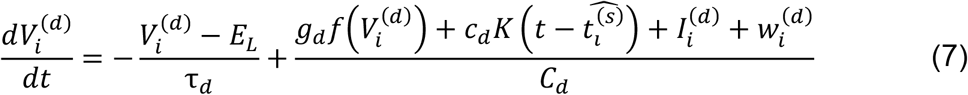

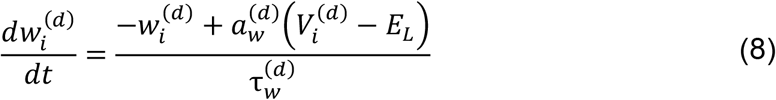

The voltages 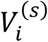 and 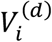 are initialized to -70mV, whereas 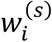 and 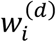 are initialized to 0. The function *f* is defined as 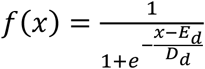 and *K* is a rectangular kernel with an amplitude of 1 between 0.5 ms and 2.5 ms. The neuron spikes when 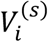 reaches *V*_T_ = −50mV, after which 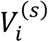 is reset to -70mV.

The input current to the somatic compartment of a landmark neuron is given by 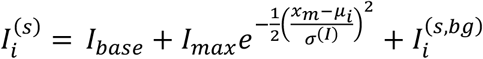. The locations *μ*_i_ where 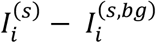 is maximized are selected from a Gaussian distribution centered at 220 cm with a standard deviation of 30 cm. The input current to the dendritic compartment of a landmark neuron is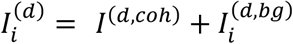. The current *I*^(*d,coh*)^ is calculated as 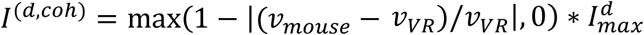. Both background currents 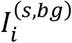 and 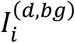 are modelled with an Ornstein-Uhlenbeck process with a mean of 0: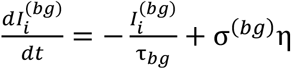, where η is white noise.

In the somatic inputs only condition (no inputs to dendrites), the somatic compartment receives an input of 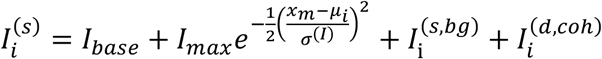 and the dendritic compartment has an input of 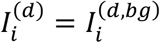.

The continuous attractor is modeled as a reduced, low-dimensional model of a line attractor (Ocko et al. 2018). Velocity inputs move the peak of the activity bump *x*_*att*_ in proportion to the mouse’s velocity *v*(*t*):

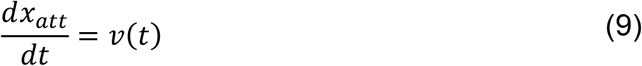

Spiking from landmark neurons can also move *x*_*att*_. Spikes from landmark neurons are convoluted with an exponential filter:

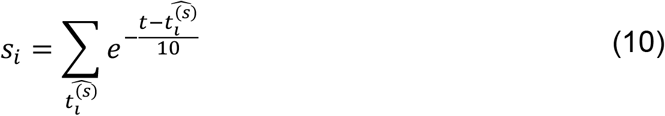

The resulting aggregated value generated by spiking multiplied by a sigmoidal function as follows:

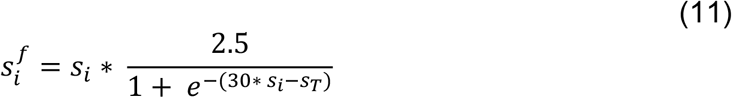

The resulting force 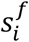 moves the activity bump moves towards the landmark cell’s receptive field center:

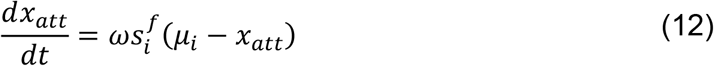

The initial location of *x*_*att*_ is determined by averaging the mouse’s starting location over the trials. ω is a normalization factor calculated as 1/number of simulated neurons. For each trial, the mouse’s velocity *v*(*t*) is determined by cutting out timesteps where the mouse’s recorded velocity exceeds 1 m/s and fitting a function to the data with linear interpolation. Unless otherwise stated, each simulation consists of 100 trials using 100 neurons.

### Single compartment model

The single compartment model was based on the 2-compartment model but without the dendritic compartment. The dynamics of the compartment are:

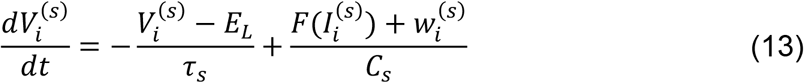

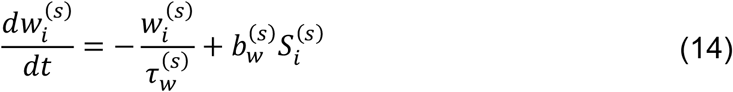

where the function *F* is defined to be

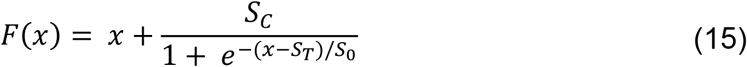

in the supralinear case, and *F*(*x*) = *x* otherwise. The somatic compartment receives an input of 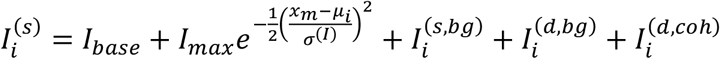.

The variables are as described in Table 1, with the addition of parameters *S*_*C*_, *S*_*T*_ and *S*_0_ describing the supralinear integration. For the simulations in supplementary figure 7, the following parameter values were used:

**Table 1:** Variable descriptions for model and simulations.

| Variable | Description |
| --- | --- |
| $V_i^{(s)}$ | Voltage of somatic compartment for the $i^{\text{th}}$ landmark cell |
| $w_i^{(s)}$ | Adaptive current for somatic compartment of the $i^{\text{th}}$ landmark cell |
| $S_i^{(s)}$ | Spike train of the $i^{\text{th}}$ landmark cell |
| $V_i^{(d)}$ | Voltage of dendritic compartment for the $i^{\text{th}}$ landmark cell |
| $\widehat{t}_i^{(s)}$ | Spike times of the $i^{\text{th}}$ landmark cell |
| $w_i^{(d)}$ | Adaptive current for dendritic compartment of the $i^{\text{th}}$ landmark cell |
| $I_i^{(s)}$ | Input current to somatic compartment of the $i^{\text{th}}$ landmark cell |
| $I_i^{(d)}$ | Input current to dendritic compartment of the $i^{\text{th}}$ landmark cell |
| $x_m$ | Mouse location |
| $\mu_i$ | Location of peak current of $i^{\text{th}}$ landmark cell |
| $I_i^{(s,bg)}$ | Background input to somatic compartment of $i^{\text{th}}$ landmark cell |
| $I_i^{(d,coh)}$ | Input to the dendritic compartment of $i^{\text{th}}$ landmark cell from the coherence signal |
| $I_i^{(d,bg)}$ | Background input to dendritic compartment of $i^{\text{th}}$ landmark cell |
| $\sigma^{(bg)}$ | Parameter for Ornstein-Uhlenbeck process for background noise |
| $x_{att}$ | Location of the peak of the steady state activity bump on the attractor |
| $v(t)$ | Mouse velocity |
| $s_i$ | Activity of $i^{\text{th}}$ landmark cell |
| $s_T$ | Threshold value for gating force from landmark neurons to attractor |
| $f_i$ | Force applied to attractor |

**Table 2:** Model parameters and set values.

| Parameter | Value |
| --- | --- |
| $E_L$ | -70 mV |
| $g_s$ | 1300 pA |
| $\tau_s$ | 16 ms |
| $C_s$ | 370 pF |
| $\tau_w^{(s)}$ | 100 ms |
| $b_w^{(s)}$ | -200 pA |
| $g_d$ | 1200 pA |
| $c_d$ | 2600 pA |
| $\tau_d$ | 7 ms |
| $C_d$ | 170 pF |
| $a_w^{(d)}$ | -13 nS |
| $\tau_w^{(d)}$ | 30 ms |
| $E_d$ | -38 mV |
| $D_d$ | 6 mV |
| $V_T$ | -50 mV |
| $I_{base}$ | 250 pA |
| $I_{max}$ | 800 pA |
| $\sigma^{(l)}$ | 5 |
| $I_{max}^d$ | 150 pA |
| $s_T$ | 1.75 a.u. |

**Table 3:** Model parameters and set values for single compartment models.

|  |  |
| --- | --- |
| $S_C$ | 2000 pA |
| $S_T$ | 1000 pA |
| $S_0$ | 10 |

## Acknowledgements

We thank Enrique Toloza and Jakob Voigts for constructive input on developing the model, and Ila Fiete, Courtney Yaeger, and Raul Mojica for helpful feedback on the manuscript. This work was supported by the NIH (RO1NS106031), a Klingenstein-Simons Fellowship, and the NEC Corporation Fund for Research in Computers and Communications at MIT.

## Author Contributions

L.F.F. performed experiments, analyzed data, designed and helped build the model, made figures, and wrote the manuscript. L.X. helped design and built the model and also analyzed model data. K.T.M. collected experimental data and assisted with analysis. M.T.H supervised all aspects of the project and helped write the manuscript.

## Data availability

All data needed to evaluate the conclusions in the paper are present in the paper and/or the Supplementary Materials. Additional data will be deposited in Zenodo upon acceptance.

## Supplementary materials

Supplementary figures 1-9.

