## Supplementory materials for "Learning to use landmarks for navigation amplifies their representation in retrosplenial cortex"

### Supplementary Materials

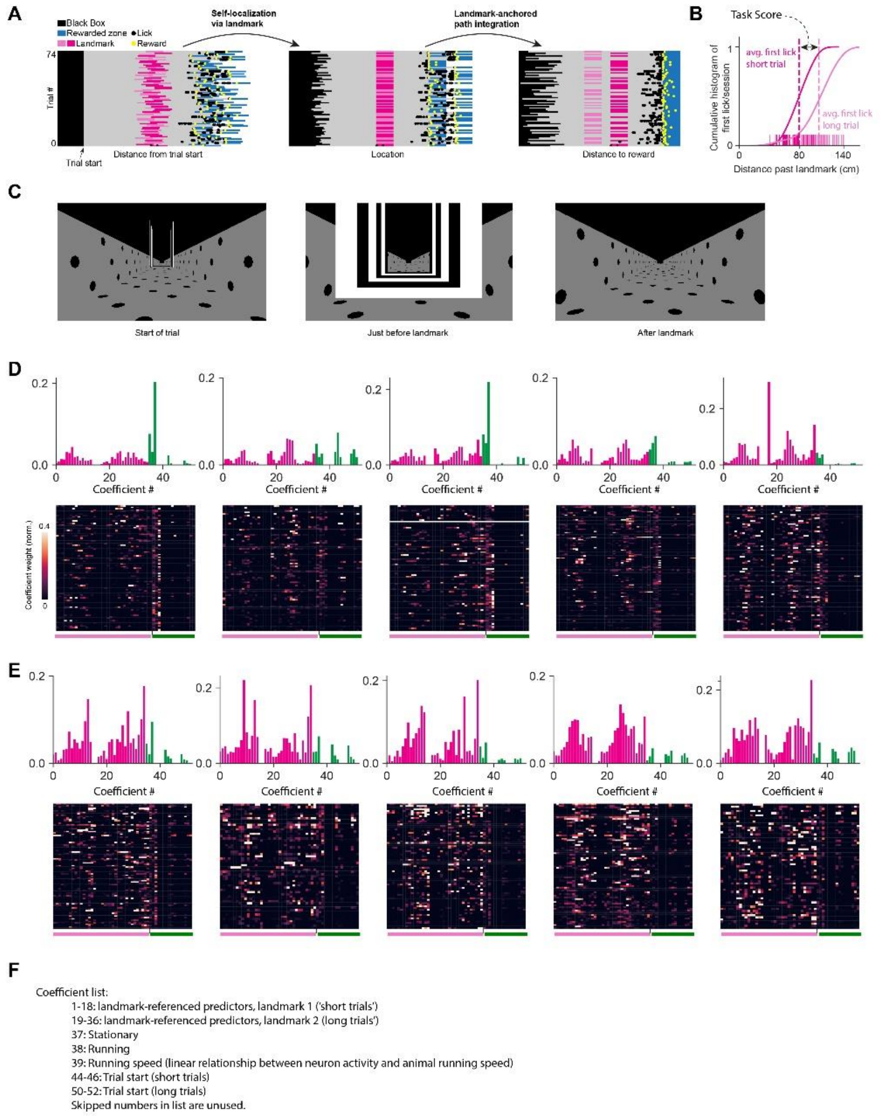


**Figure S1: GLM coefficients in individual animals.** (**A**) Schematic of the spatial computations required to successfully execute out the task. The three panels show behavior in an example session. The color code indicates the different sections of the track and each line represents one trial. As an animal progresses through a trial, it starts without yet knowing where it is relative to the landmark and respective reward (left). As it approaches the landmark it can localize itself relative to the landmark (middle) and then, by licking in the right location (black and yellow dots), it can trigger reward release (right). (**B**) Cumulative histograms of first lick locations after the landmark for example an example session. Dashes along the bottom indicate individual first licks/trial. Pink shading indicates which landmark was shown. Dashed lines indicate average first lick location on short and long trials. (**C**) Mouse field-of-view at three points along Track 1. (**D**) Histograms (top) and coefficient heatmaps (bottom) for n=5 mice included in this study before learning the task (Task score < 40 cm). (**E**) same as in (D) but for proficient animals (task score > 40 cm). (**F**) Full list of predictors for GLM.

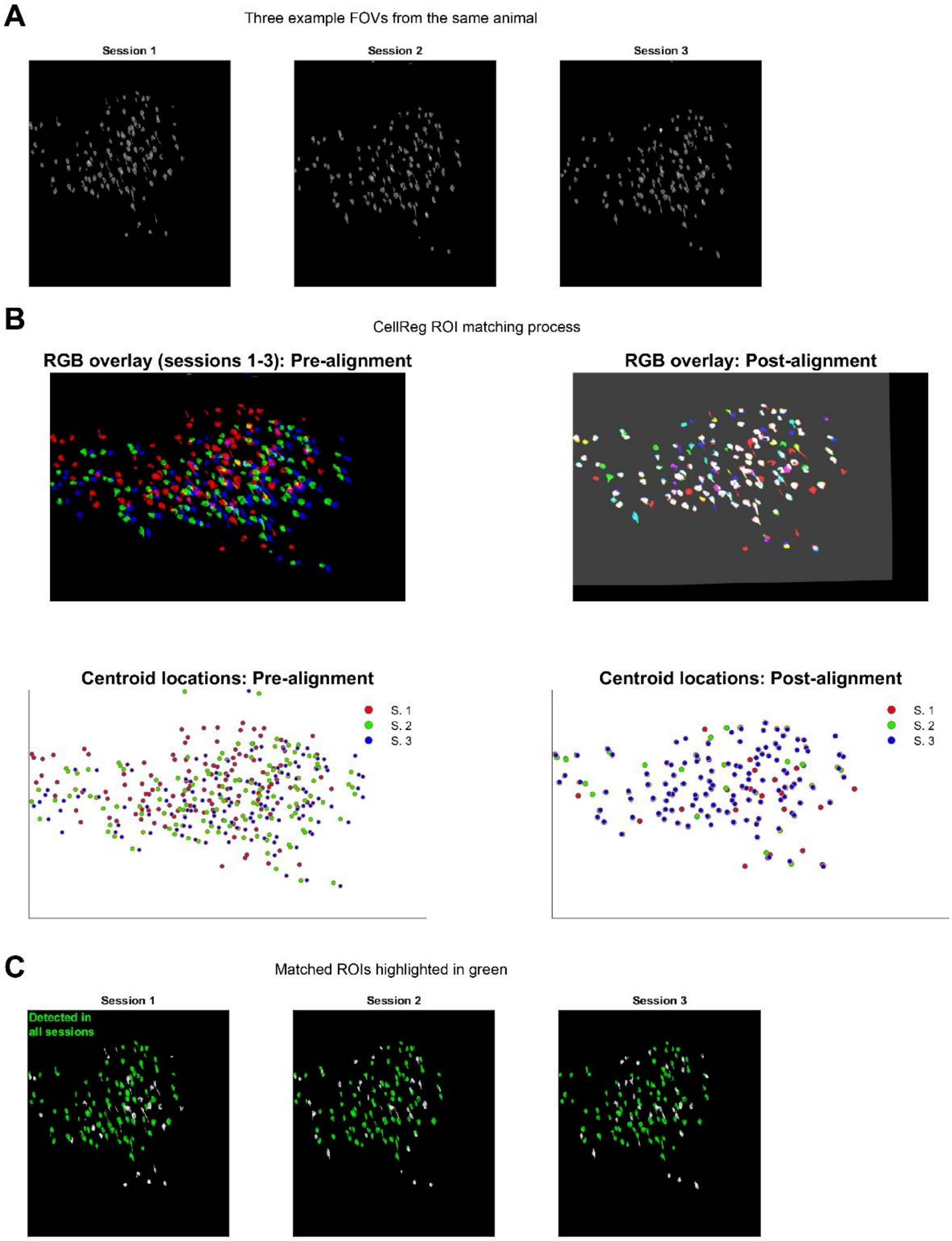


**Figure S2: Registering ROIs across multiple recording sessions.** (**A**) Detected ROIs in example field of view in three consecutive recording sessions. (**B**) CellReg registration of three sessions (red, green and blue). Left: detected ROIs (tops) and ROI centroid locations (bottom) of an example FOV. Right: ROIs are shifted to match ROI locations across sessions. Unmatched ROIs were not used for cross-day analysis. (**C**) Result of matching ROIs across days. Matched ROIs are highlighted in green.

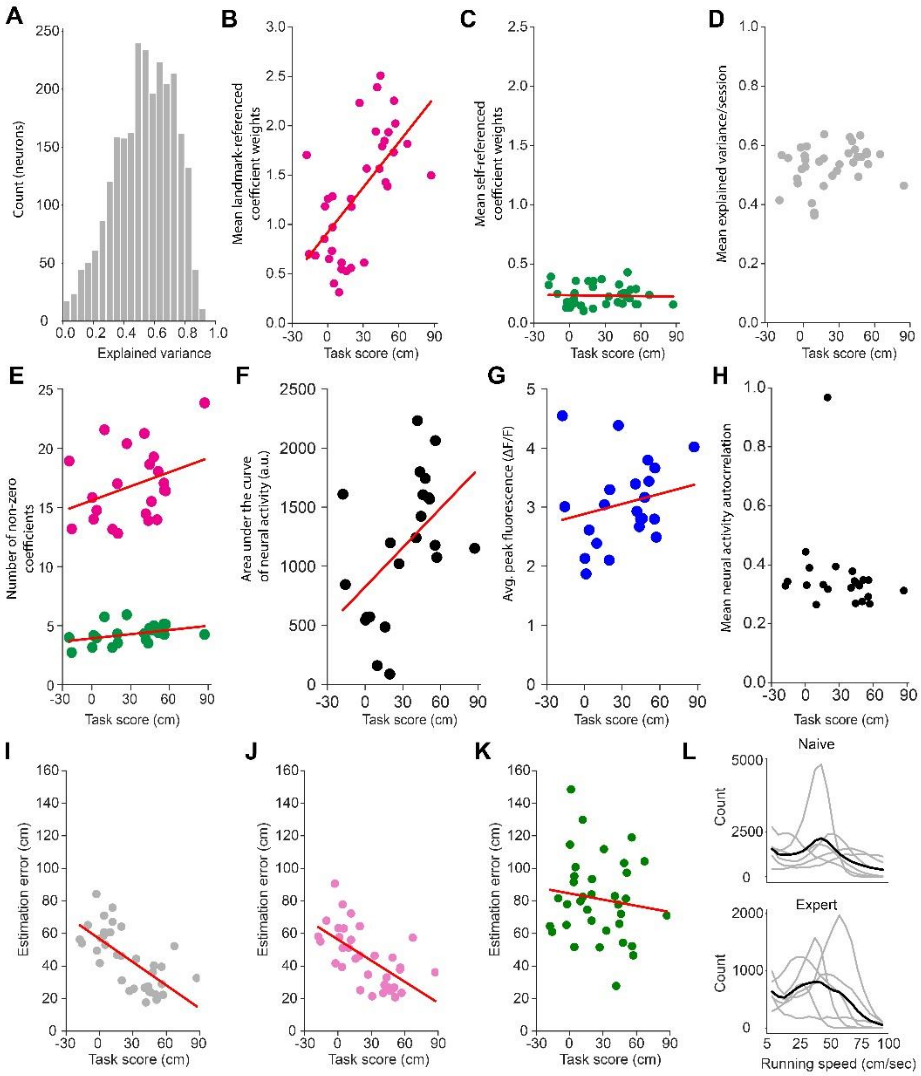


**Figure S3: GLM coefficients, $r^2$, and location reconstruction as a function of task score.** (**A**) Explained variance ($r^2$) by GLM for all recorded neurons in all sessions. (**B**) Landmark-referenced coefficient weights (mean of sum of all landmark-referenced coefficients) for all neurons in all recorded sessions (n=35 sessions, 5 animals, mean ± SEM: 75.51 ± 3 neurons/session, Spearman rank correlation: ρ=0.618, p<0.001). (**C**) Same as (B) but for self-referenced coefficient weights (Spearman rank correlation: ρ=0.014, p=0.936). (**D**) Explained variance (mean of all neurons in a given session, Spearman rank correlation: ρ=0.275, p=0.1). (**E**) Number of non-zero GLM coefficients as a function of task score. Pink: landmark-referenced coefficients, green: self-referenced coefficients. Neither are significantly correlated with task score. (**F**) Neural activity measured as mean area under the curve for all tracked neurons in a given session as a function of task score. A significant correlation between tasks core and AUC exists (Spearman rank correlation: ρ=0.469, p=0.028). (**G**) Same as (F) but instead of area under the curve, the peak fluorescence is measured. No significant correlation exists between mean peak fluorescence and task score. (**H**) Mean autocorrelation of all neurons for all tracked sessions showing no significant correlation between autocorrelation and task score. (**I**) Position estimation error of a Bayesian decoder using activity predicted by the GLM. (**J**) Same as (I) but for activity predicted solely by landmark-referenced coefficients. (**K**) Same as (J) but for activity predicted solely by self-referenced coefficients. (**L**) Running speed histograms for n=5 animals in naïve and expert (task score > 20 cm) condition.

A

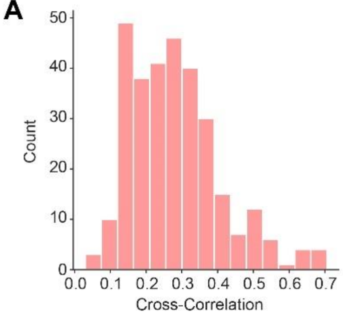


**Figure S4**: **Distribution of cross-correlation values of neural activity across session**. (**A**) Histogram of cross correlation values for neurons tracked across expert session 1-3 as shown in Fig. 2 (Mean ± SEM: 0.28 ± 0.007).

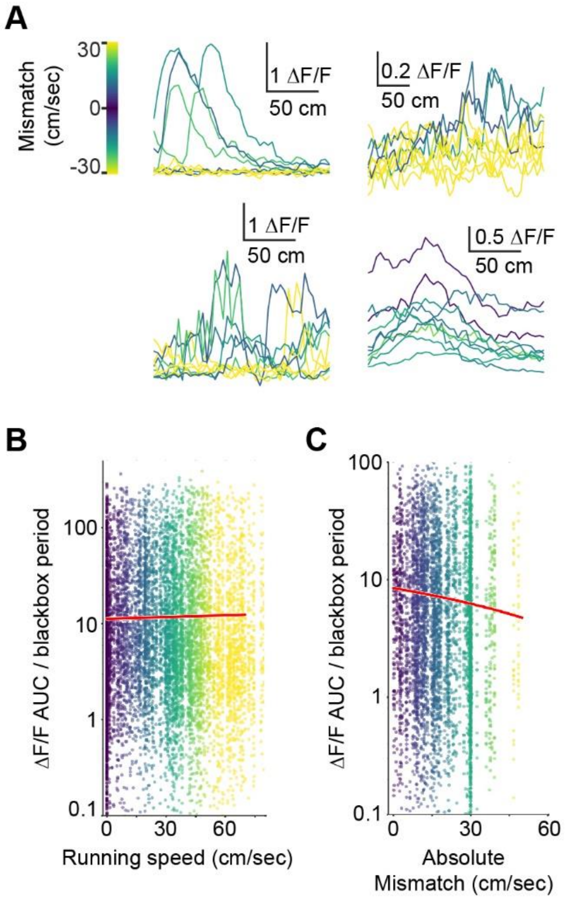


**Figure S5: Visuo-motor mismatch responses.** (**A**) Overlaid single trial responses of different degrees of mismatch from four example neurons with different response patterns. (**B**) Visuo-motor mismatch responses in between trials when animals were placed in a black box for three seconds. No significant correlation exists between mismatch and neural responses (Spearman $\rho$=-0.0055, p=0.52). (**C**) Visuo-motor mismatch responses during virtual navigation without linear speed correction (Spearman $\rho$=-0.108, p<0.001).

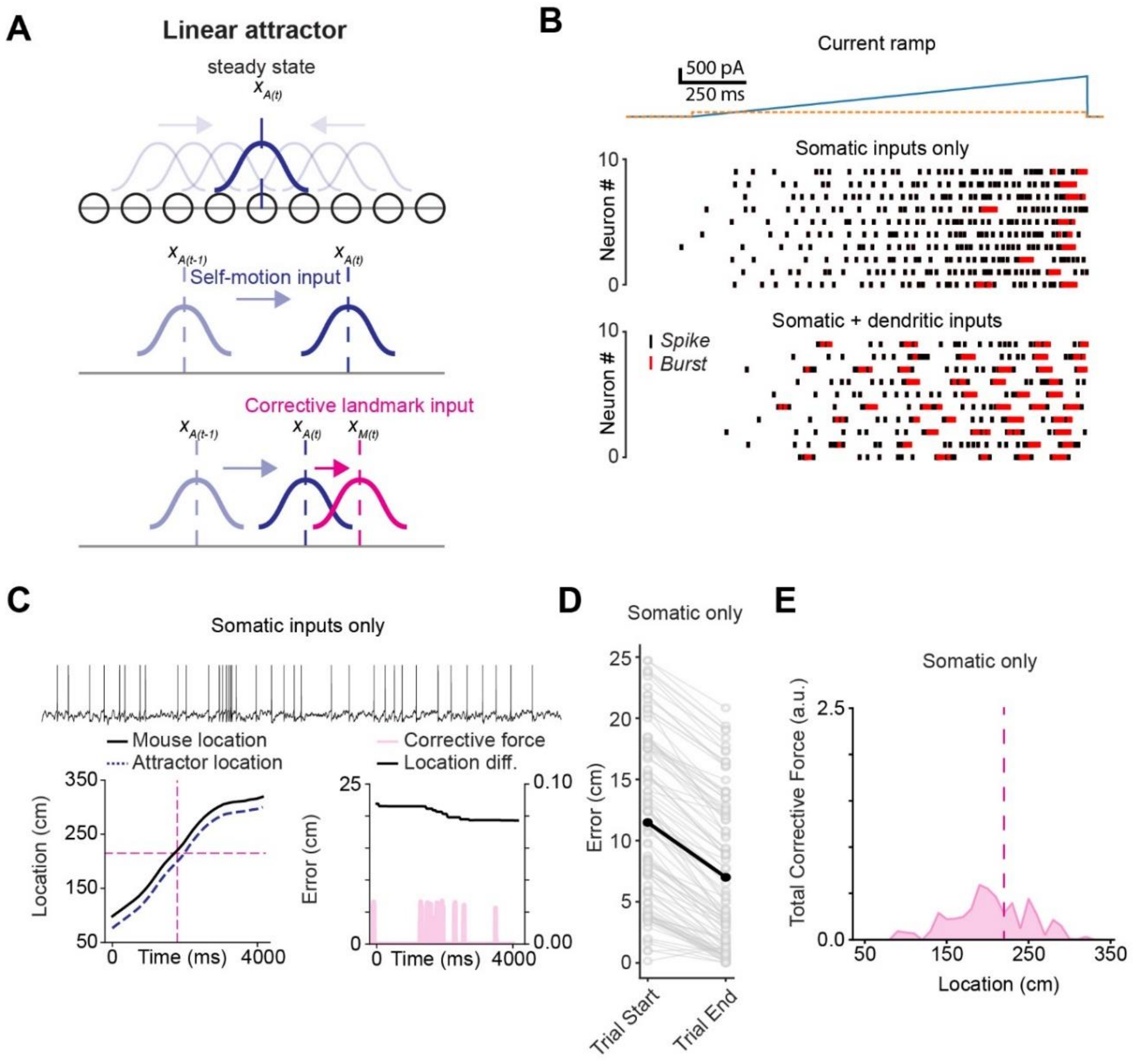


**Figure S6: Attractor schematic, burst probability, and simulated trials showing self-localization performance in the absence of dendritic inputs.** (**A**) Schematic of the linear attractor. Agent location ($X_A$ at time t) is represented as a Gaussian activity bump centered on the self-localization estimate (top). Self-motion feedback moves the activity bump forward (middle). Landmark inputs to the attractor network shift the activity bump towards the center of the landmark neuron's receptive field center XM (bottom). (**B**) Current ramp input in the presence (bottom) and absence (middle) of concurrent dendritic inputs. Top: current injection amplitude. Orange indicates dendritic, blue indicates somatic injection. In the 'Somatic inputs only' condition both currents are injected into the soma compartment. (**C**) Top: example neuron trace. *Bottom left*: Actual animal location and self-localization estimate of attractor in one trial. *Bottom right*: Self-localization error and corrective force. (**D**) Error at the beginning and at the end of 100 simulated trials (Mean final error: 7.0 ± 0.61 cm. Paired t-test: $p < 0.001$). (**E**) Mean corrective force as a function location (mean corrective force AUC ± SEM: 52.8 ± 1.50)

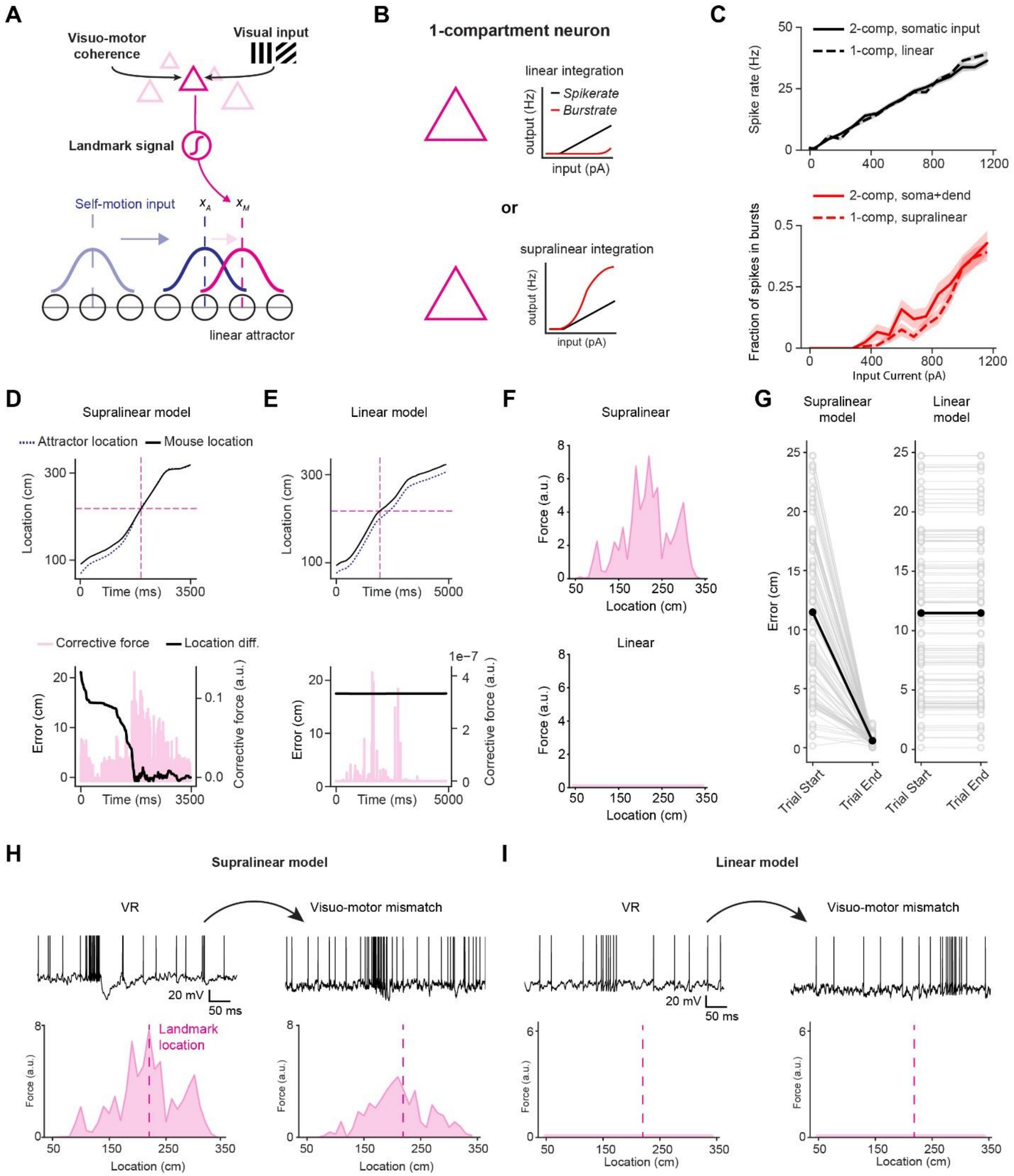


**Figure S7: Models with single-compartment neurons do not exhibit behavioral context-dependence.** (**A**) Schematic of the full model using single-compartment neurons. Two models with the same architecture but with either linear or supralinear single-compartment neurons were used. (**B**) Schematics of linear and supralinear model neurons. (**C**) Comparison of spike rate and fraction of neurons in bursts for linear and supralinear single-compartment neurons, as well as two-compartment neurons, in response to current input. (**D**) Example trial of model with supralinear single-compartment neurons. Top: Mouse location vs. attractor location, bottom: location error of attractor and force exerted by cortical neurons. (**E**) Same as (D) but with model using linear single-compartment neurons. (**F**) Force exerted on the attractor as a function of location. (**G**) Self-localization error at the start and end of trials. (**H**) Model using supralinear single-compartment neurons during virtual navigation and visuo-motor mismatch. Top: example neuron trace. Bottom: total force exerted on the attractor as a function of location. (**I**) Same as (H) but for model using linear neurons.

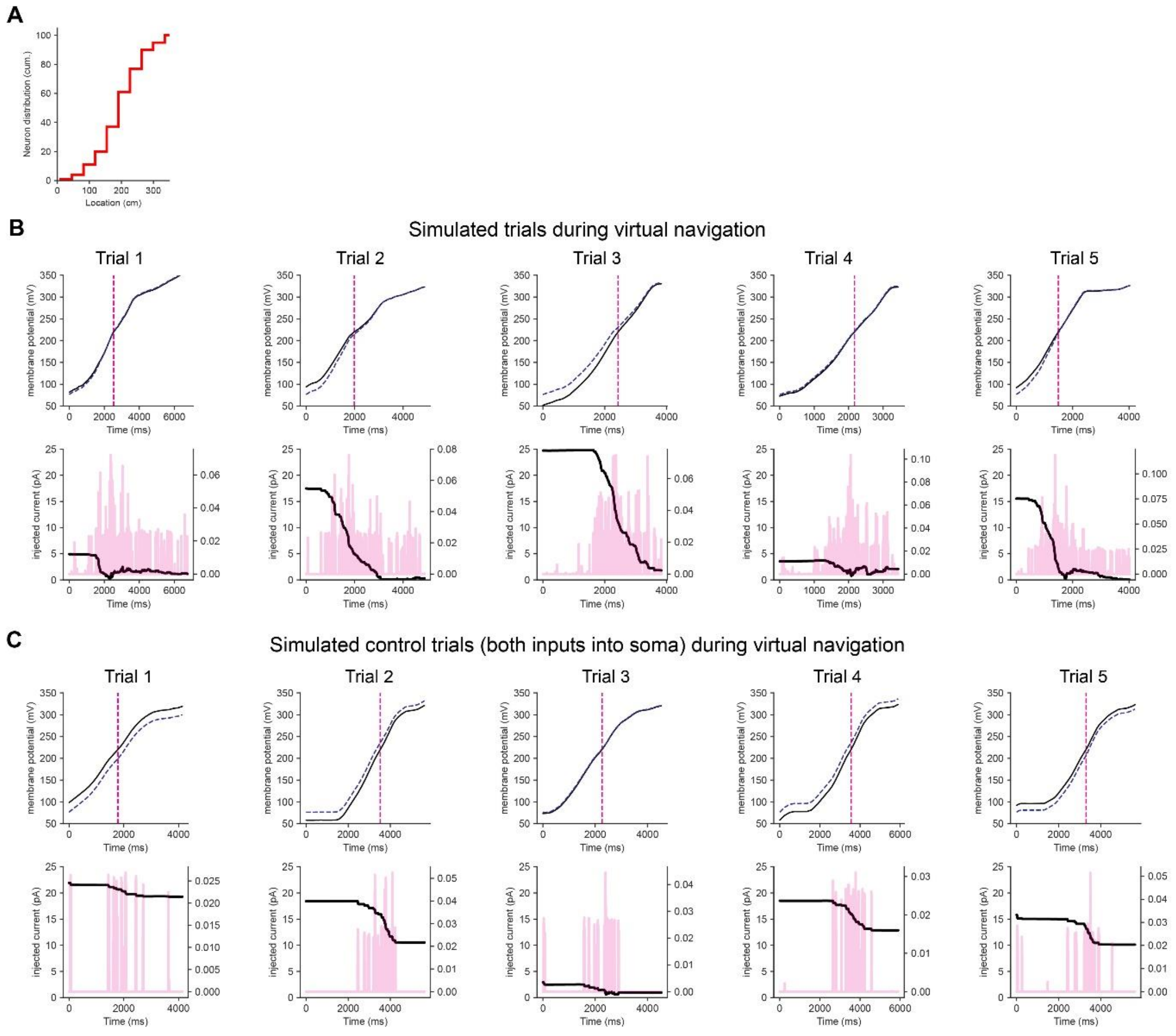


**Figure S8: Landmark receptive field distribution and example simulated trials of 2-compartment neurons.** (**A**) Cumulative distribution of visual receptive field center for multicompartment neurons during simulations. (**B**) Example simulated trials during virtual navigation. Top: Mouse (black) and attractor (blue dashed) location. Bottom: Difference between mouse and attractor location (black) and force exerted on the attractor (pink) (**C**) Same is (B) but for simulated trials during control condition in which both current inputs were injected into the soma.

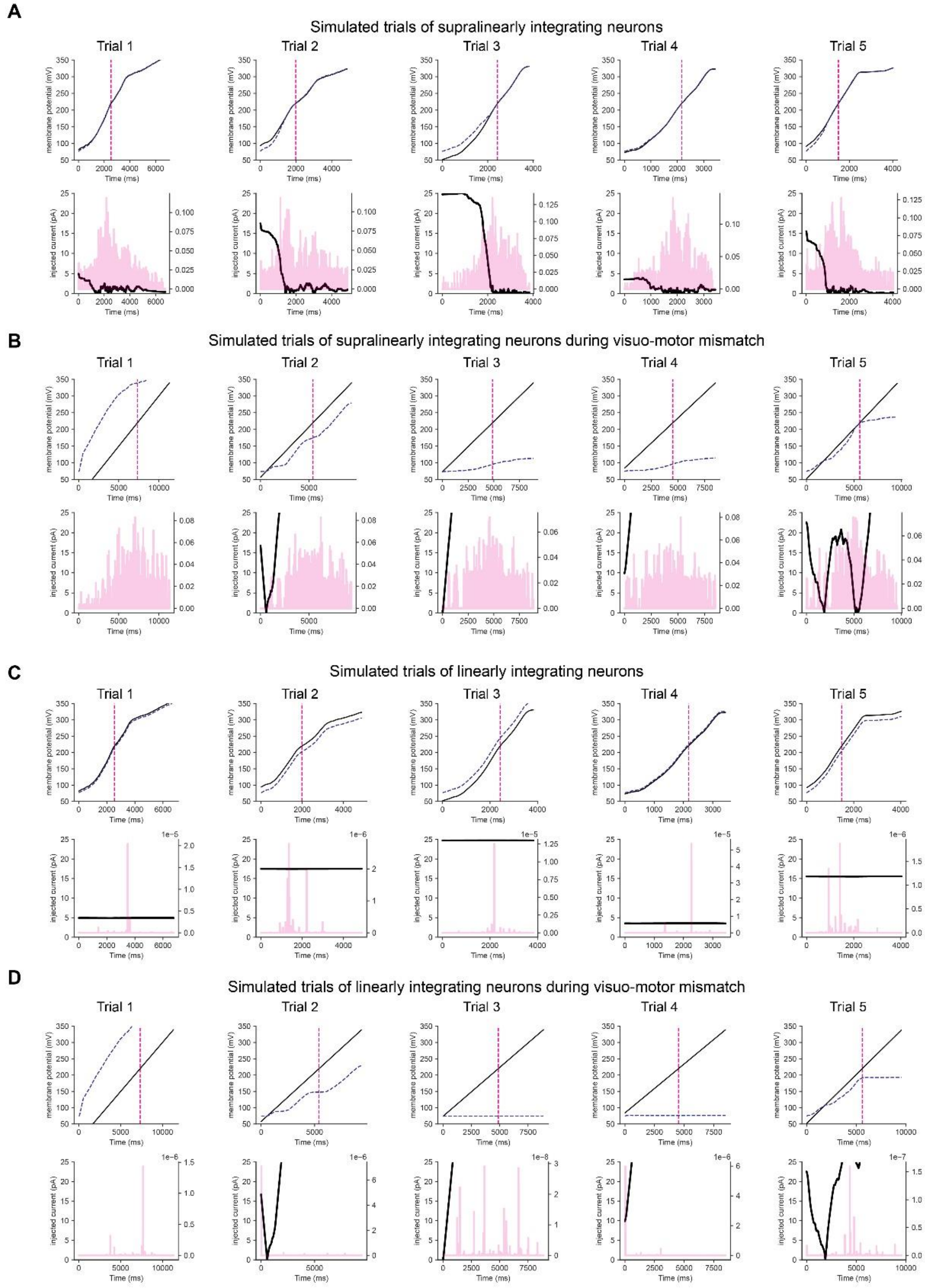


**Figure S9: Example simulated trials of single-compartment neurons** (**A**) Example trials for supralinearly integrating neurons. Top: Mouse (black) and attractor (blue dashed) location. Bottom: Difference between mouse and attractor location (black) and force exerted on the attractor (pink) (**B**) Same as (A) but during visuo-motor mismatch. (**C,D**) Same as (A,B) bur for linearly integrating neurons.
